# Naturalistic Movies and Encoding Analysis Redefine Areal Borders in Primate Visual Cortex

**DOI:** 10.1101/2023.12.25.573299

**Authors:** Daisuke Shimaoka, Yan Tat Wong, Marcello GP Rosa, Nicholas Seow Chiang Price

## Abstract

Accurate definition of the borders of cortical visual areas is essential for the study of neuronal processes leading to perception. However, data used for definition of areal boundaries has suffered from issues related to resolution, uniform coverage, or suitability for objective analysis, leading to ambiguity. Here, we present a novel approach that combines widefield optical imaging, presentation of naturalistic movies, and encoding model analysis, to objectively define borders in the primate extrastriate cortex. We applied this method to test conflicting hypotheses about the third-tier visual cortex, where areal boundaries have remained controversial. The results support a hypothesis whereby an area contains representations of both the upper and lower contralateral quadrants (DM) is located immediate anterior to V2, and unveil pronounced tuning preferences in the third-tier areas. High-density electrophysiological recordings with a Neuropixels probe confirm these findings. Our encoding-model approach offers a powerful, objective way to disambiguate areal boundaries.

## Introduction

While cortical areas are generally perceived as distinct modules both functionally and anatomically, the actual situation is often more nuanced than commonly believed. For example, in the visual cortex, there is widespread consensus regarding the functionally-^1^ and anatomically^2^ defined boundaries and organization of early stages of visual processing, including primary (V1) and secondary (V2) visual areas, and the middle temporal (MT) visual areas. These areas contain neurons that exhibit well-defined constellations of response properties, form complete maps of the contralateral visual field, and coincide with specific architectural patterns, and form connections with other cortical areas and subcortical nuclei, which are characterized by specific laminar patterns.

However, the borders of areas remain unsettled in some of higher-order cortical areas. For instance, the precise configuration of the third-tier visual cortex, situated immediately anterior to V2, remains controversial^3,4^. The seemingly simple issue of defining areal boundaries in a relatively early stage of hierarchical visual processing remain unresolved, in part, because histologically-defined boundaries have historically been drawn based on subjective assessment of anatomical connections that often vary according to the part of the visual field being represented^5–7^. Greater objectivity may be afforded by cellular-resolution techniques such as extracellular unit recordings with microelectrodes to map receptive fields of single neurons. However, the use of microelectrodes also has limitations, as it encompasses a limited portion of the cortex at a time. In fact, it if often the case that choices have to be made in experimental design between covering large areas, or studying the physiological properties of neurons in detail. Even the latest Neuropixels 2.0 array covers 10 x 1mm area of the cortex. An ideal method for testing competing hypotheses of areal boundaries therefore requires receptive field mapping from large portions of cortex. Moreover, the existing approaches require *a priori* selection of the battery of parameters to be tested, which may not be those relevant to reveal differences between areas.

Widefield imaging is a promising candidate for objectively defining areal borders, as it records functional responses from large portions of uninterrupted cortex simultaneously. However, previous imaging studies for defining areal borders, often employed synthetic stimuli such as oriented bars and dots, resulted in strong activation in V1 and V2, but not areas beyond. To overcome these limitations, in the present study, we used naturalistic movies encompassing a range of spatial and temporal frequencies, orientations, and complex visual features. To obtain retinotopic information, we applied an encoding model to characterize how neurons within each pixel of our widefield imaging data respond to visual stimuli^8^. Without needing to specify exact parameters for stimulating those areas in advance, we were able to obtain receptive fields for each camera pixel, which then can be used to sample receptive fields of any location in the imaged cortex. This approach, recently used to explore encoding of abstract modalities, has not been used for principled hypothesis testing for defining areal borders.

Here we demonstrate the application of encoding models to data obtained with widefield optical imaging is deployable to disambiguate between conflicting hypothetical organizations of the third-tier cortex (the region located immediately anterior to V2^9^) in new world primates. This approach enabled us to objectively compare two prominent hypotheses on the organization of this region, based on high-resolution retinotopic maps. Our results reveal the presence of a third-tier region that represents the upper visual field immediately anterior to V2, and S-shaped receptive field transitions between regions coding the upper and lower visual fields – both unambiguous predictions of the hypothesis whereby a dorsomedial area (DM) is located in this region, instead of a continuous third visual area that adjoins most of V2^10,11^. These observations were confirmed with high-density electrophysiological recordings with a Neuropixels probe. The encoding model approach provides additional insight into tuning properties beyond receptive fields, showing marked preferences for high temporal frequencies and speeds. These observations provide resolution to one of the most enduring controversies about the primate extrastriate cortex, and demonstrate the powerful capability of our encoding-model approach in defining and characterizing individual areas.

## Results

To examine the functional organization of the third-tier visual areas with respect to neighboring visual areas, we conducted widefield intrinsic optical imaging of the dorsal-posterior portion of the cortex, encompassing multiple visual areas including V1, V2, third-tier visual areas and posterior-parietal cortex (**Figure 1A-C**). We analyzed the retinotopic organization of the population receptive fields corresponding to every pixel within the cranial window. To estimate receptive fields during presentation of naturalistic movies, we employed a motion-energy model, developed to describe population neuronal activities of early visual cortical areas^12,13^. The model is composed of Gabor filters (kernels) distributed across the visual field, with varying spatio-temporal frequencies, sizes, orientations, and directions. We fit all the kernels in the model with the observed imaging signal in each camera pixel. An example of this fitting process for a single pixel at the center of the cranial window (**Figure 1D**) is shown in **Figure 1E-I**. The fitted model captured the signal fluctuation, with 0.33 correlation with the observed signal (**Figure 1E, F**). The distribution of the estimated 6555 kernel coefficients exhibited a skewed pattern (**Figure 1G**). The 4 kernels with highest coefficients shared the same position in the visual field, and 3 had the same orientation, but a range of spatial frequencies was observed (**Figure 1H**). From the estimated coefficients, we obtained the receptive field and its center in the visual field (**Figure 1I**).

**Figure 1.**
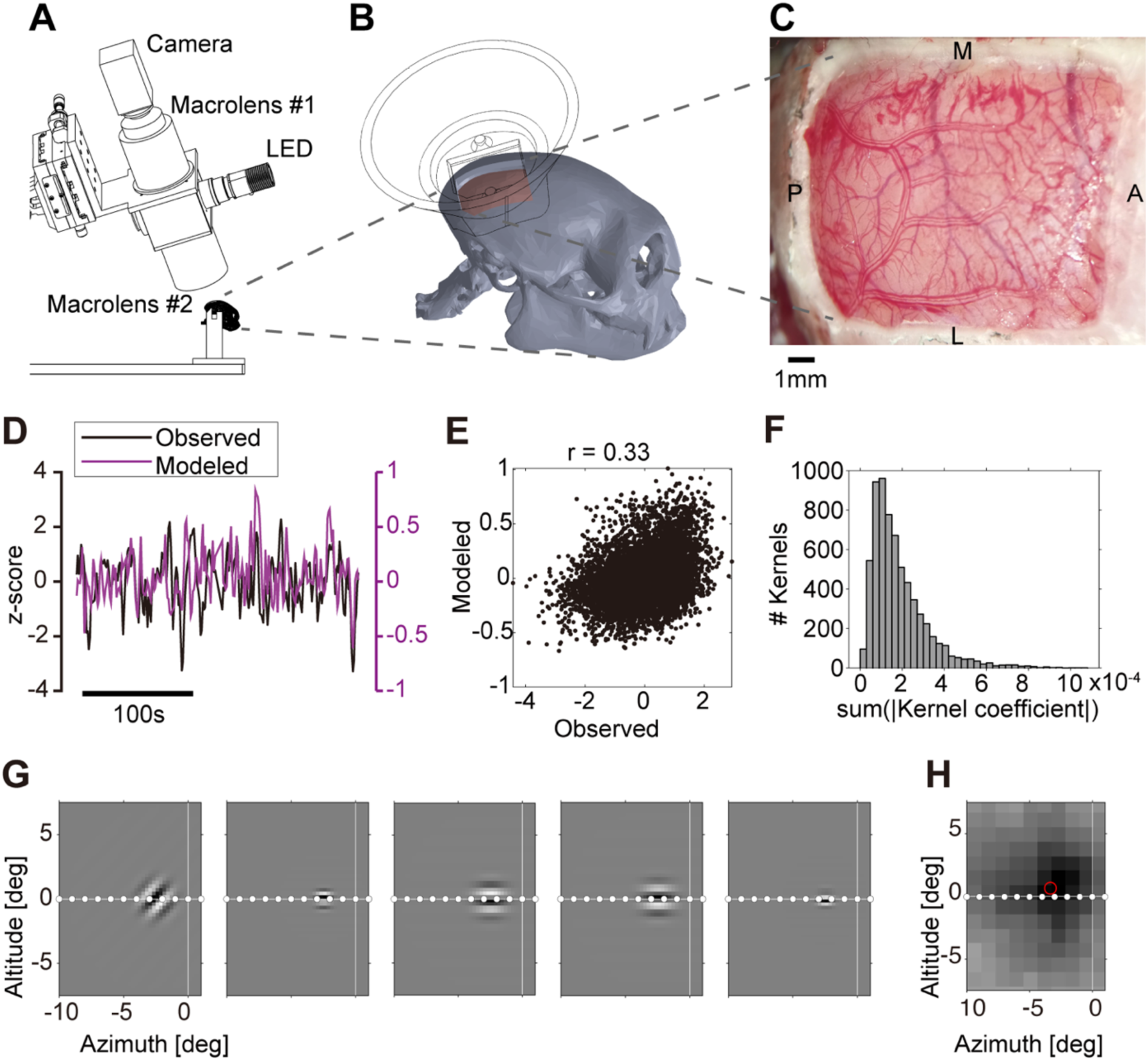
Method for high-resolution receptive field mapping. **A-D.** Widefield intrinsic optical imaging. **A.** Green light (LED) was applied to the cortex, and the reflected optical signals passed through two macro lenses before capture by a CCD camera. **B.** To ensure optical isolation from ambient light and minimize pulsation artifacts, a custom 3D-printed cone (shown as a wireframe) was placed over the exposed cortex. The cone’s lid was fitted with a glass coverslip. **C**. The exposed cortical region, observed from above, covered 10×13 mm. **D**. The observed imaging signal (black) and its corresponding modeled signal (purple), in one pixel. The modeled signal is obtained through fitting the motion-energy model incorporating 6555 linear spatio-temporal filters. **E.** The modelled signal (ordinate) exhibits a substantial correlation with the observed signal (abscissa). **F.** The distribution of coefficients for the linear filter is presented. These coefficients are estimated from the observed imaging signal, summed over the time axis of the filter. **G.** Linear filters with highest coefficients from panel **F** are highlighted. Their representation within the visual field is depicted. Notably all the five filters encode visual field approximately 4 degrees left of the fovea. **H.** The receptive field’s position for the model, employing all the fitted filter coefficients from panel **F** is depicted. The peak location is designated as the red circle. This receptive field was obtained through *in silico* simulation of the model exposed to sparse noise stimuli.

### Hypothesis-driven testing of areal organization of third-tier areas

We applied the encoding model approach to investigate cortical organization in and around the third-tier visual areas - whose precise organization remains elusive, particularly regarding how many areas form the dorsal visual cortex immediately rostral to V2, and where their boundaries lie^14,15^. The single third-tier area hypothesis, illustrated for marmosets, (**Figure 2A**) proposes the presence of an elongated third visual area, immediately anterior to V2 and covering the entire visual field^4,16^. The dorsal segment of the third-tier area exclusively processes the lower visual field and at its posterior border, it interfaces a separate visual area (DM) that includes a region encoding the upper visual field.

**Figure 2.**
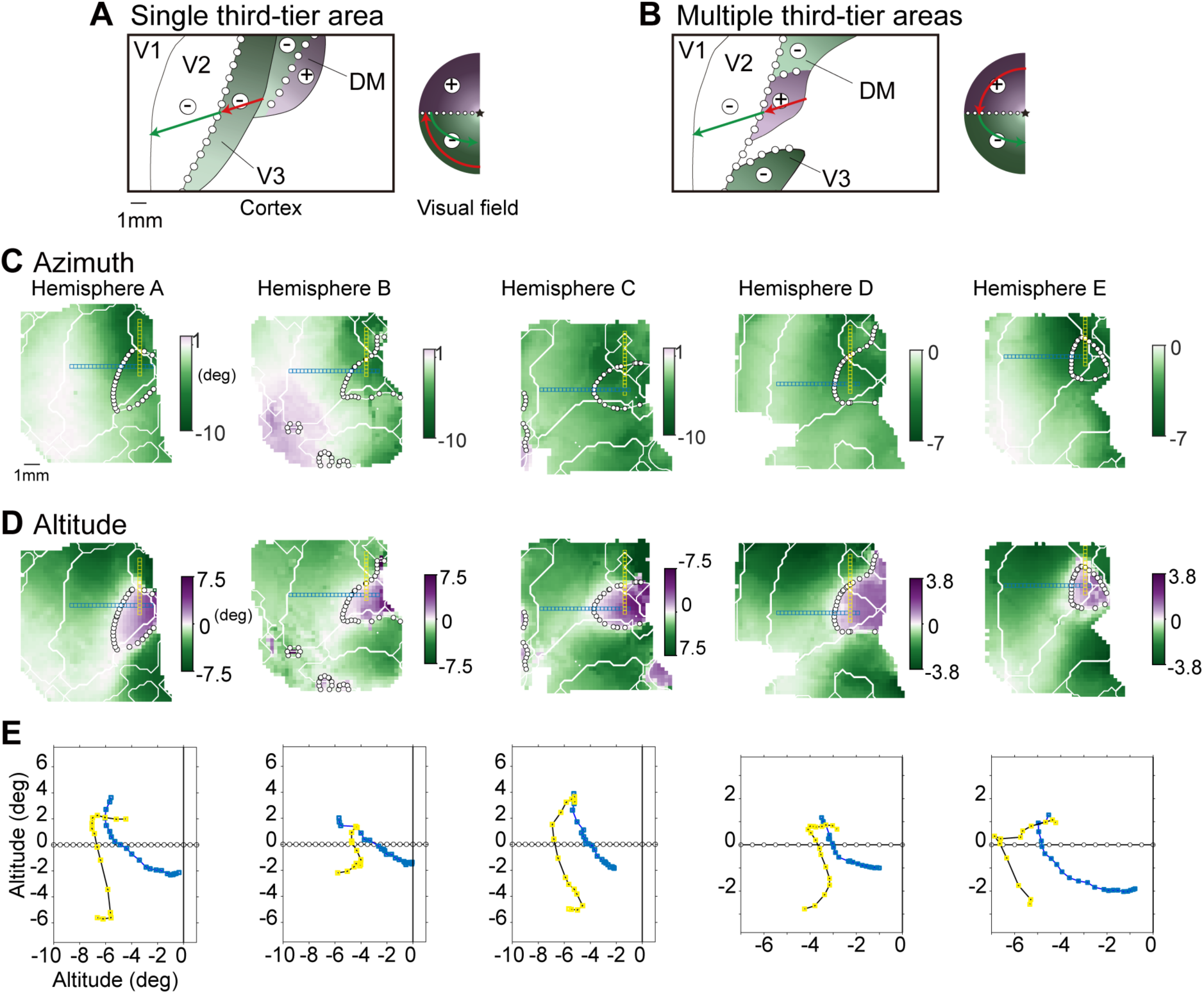
Hypothesized and Measured Areal Organization of Third-Tier Visual Areas. **A**. Single area hypothesis. The region immediately rostral to the dorsal portion of V2 contains a solitary third-tier visual area, V3(d). The two regions are demarcated by the horizontal meridian (represented by a line with open circles), while V2 and V1 is separated by the vertical meridian. The cortical positions indicated by the red and green arrows in the left panel correspond to specific receptive field locations within the visual field. The distance from the fovea (star symbol) is represented by a light-gray shading. This hypothesis predicts that the dorsal portion of V3 covers only lower visual field (-) but not upper visual field (+). **B**. Two-areas hypothesis. This hypothesis proposes that the region right rostral to V2 comprises two third-tier visual areas (V3 and DM). This hypothesis predicts a limited region (encompassed by the red arrow) in the dorsal part of the cortex encodes the upper visual field. **C, D.** Altitude and azimuth of the preferred visual field positions are mapped onto the imaged cortical region. Only pixels with successful model fitting are shown. The preferred positions are calculated from the peak of the receptive field in each camera pixel. Solid lines indicate estimated boundaries of cortical areas, computed with the map of visual field sign (Figure 3). Dotted lines signify vertical meridian, derived from the preferred altitude. Notably in panels **C**, a portion of the imaged cortical region encodes the upper visual field, as highlighted in purple. **E.** A sequence of receptive field locations associated with pixels shown as blue and yellow squares in panels **C**&**D**. (**A** and **B**, adopted and modified from Rosa and Tweedale (2013))

Conversely, the multiple third-tier areas hypothesis (**Figure 2B**) posits that the third-tier visual cortex comprises multiple distinct areas, with a dorsomedial area (DM in marmoset and owl monkey, V3a in Macaque) representing both the upper and lower visual fields^17,18^. Notably, this area displays an intriguing characteristic: it exhibits mirror and non-mirror representations of the visual field between regions encoding the upper and lower visual fields. The two contrasting views regarding the organization of the third-tier areas are widely observed across human and non-human primates^2,4,19,20^.

To examine these possibilities, we obtained high-resolution retinotopy maps, through application of the modeling procedure (**Figure 1**) to find the receptive field center for every pixel within the camera’s field of view. The altitude and azimuth of these receptive fields from five hemispheres imaged in four animals are represented as color-coded maps overlayed on images of the cortical surface (**Figure 2C, D**). These maps illustrate gradual transition of the preferred positions across different regions of the visual cortical areas, providing clear evidence of a well-organized retinotopic arrangement. Notably, as seen in **Figure 2C**, a specific section of the cortex (highlighted in purple) is responsible for encoding the upper visual field.

The maps of preferred altitude and azimuth let us sample receptive field positions from any part of the imaged cortex, making it possible to directly test the two hypotheses. As one such example, we examined how the RF positions in visual field travels as we virtually moved the recording sites from V2 to the region anterior to V2 (**Figure 2C-E, *blue squares***). In all five hemispheres, we found that the receptive fields move from the lower visual quadrant, cross the horizontal meridian, and reach the upper visual quadrant.

We also examined how the RF positions travel as we virtually moved the recording sites from the region encoding upper visual field to its medial region (**Figure 2C-E, *yellow squares***). In hemispheres A-D, we found a characteristic S-shaped trajectory that encompasses the upper and lower visual quadrants. This unusual trajectory with the third-tier areas was predicted by a model that balanced two constraints: smoothness in the representation within an area and congruence between adjacent areas^11^. Hemisphere E does not have sufficient coverage in the medial areas encoding the lower visual quadrant to observe this S-shaped trajectory.

Together, our approach yielded high-resolution retinotopic maps which let us examine the hypothesis-driven testing of third-tier areas. This examination revealed upper visual field representation, as predicted by the multiple third-tier areas hypothesis.

### Visual field sign

The fact that the upper field representation is immediately adjacent to a large island of lower field representation does not in itself prove adjacency between V2 and DM. To disambiguate between the hypotheses, we also need to know the locations of the representations of vertical and horizontal meridians, or more generally the field sign transitions. Visual field sign was calculated from the maps of population receptive fields (**Figure 3A**). Further, the borders between regions with positive and negative field signs were used to delineate the boundaries of visual cortical areas (white lines in **Figure 3A**). In all hemispheres, a conspicuous border between negative and positive field signs was consistently observed near the posterior edge of the cranial window, aligning with the boundary between V1 and V2. Moreover, all five maps of field sign exhibited a triangular bulge of positive field sign with anterior borders flanked by two separate regions characterized by negative sign. This bulge region was directly connected to V2, and between these two regions, a horizontal meridian (white circles) was evident.

**Figure 3.**
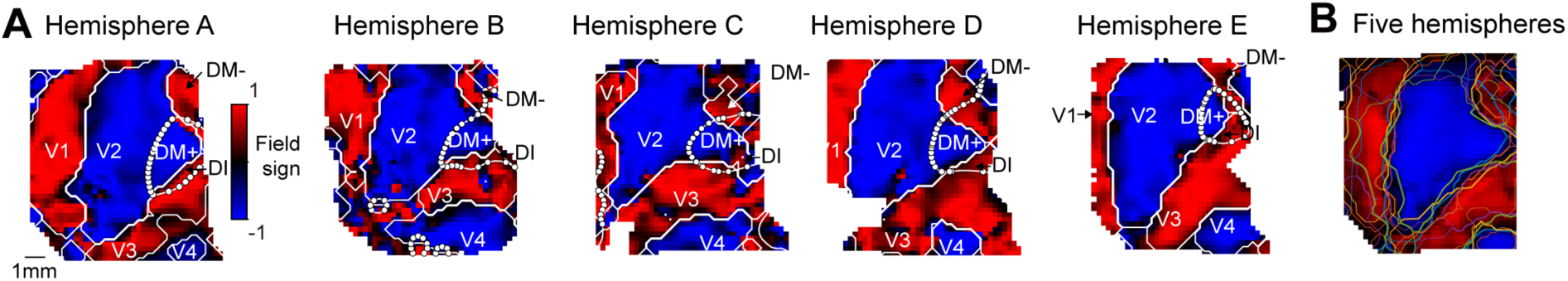
Visual Cortical Area Boundaries across Five Hemispheres. **A.** Visual field sign, computed utilizing both preferred altitude and azimuth (**Figure 2C, D**), in each of the five hemispheres. Solid lines indicate estimated boundaries of cortical areas, computed with the map of visual field sign. Dotted lines signify vertical meridian, derived from the preferred altitude (Figure 2C). **B.** A visual field sign map, averaged across the five hemispheres is represented, after aligning each map to the map from hemisphere A. Colored lines indicate estimated boundaries from each of five hemispheres.

To examine the common features of visual field sign across hemispheres, we used an image registration process to average the visual field sign map across hemispheres (**Figure 3B**). This involved aligning field sign maps for Hemispheres B-E with the map of Hemisphere A through rigid registration. The resulting averaged visual field sign map clearly retains the cross-hemisphere features articulated above. These observations indicate that the region encoding upper visual field is located immediate anterior to V2, in all five hemispheres studied.

### Imaging and Cellular Electrophysiology

The retinotopy and visual field sign maps consistently showed a region encoding upper visual field, immediate anterior to V2. However, a narrow strip may in principle exist between the two regions, that is not detectable with the limited spatial resolution of widefield imaging. Therefore, we conducted multi-unit electrophysiological recording in a subset of the experiments. Following the imaging session for Hemisphere A (**Figures 1–3**), we inserted a high-density Neuropixels probe extending for 10 mm, at an angle near tangential to the cortical surface (**Figure 4A**). The recording channels encompassed cortical regions from which the intrinsic optical signal indicated receptive fields in both upper and lower visual fields (**Figure 4C**). The spiking activity from units numbered from 1 to 6 exhibited clear spatial preferences in their responses to sparse noise stimuli (**Figure 4D**). Notably, units 5 and 6 displayed a preference for the upper visual field, corresponding with the intrinsic signal from the corresponding pixels (highlighted in red in **Figure 4C**). This alignment provides independent confirmation of the presence of the region dedicated to encoding the upper visual field. To further assess the congruence between the results obtained from the two recording modalities, we compared imaging and electrophysiology results in the preferred altitude, a key metric for testing the two hypotheses. These two recording modalities were strongly correlated (r = 0.85). Two additional penetrations, targeting more medial and lateral sites, yielded only lower visual fields. Taken together, these analyses provide compelling evidence for the existence of a cortical region where cellular spiking activity distinctly favors the upper visual field.

**Figure 4.**
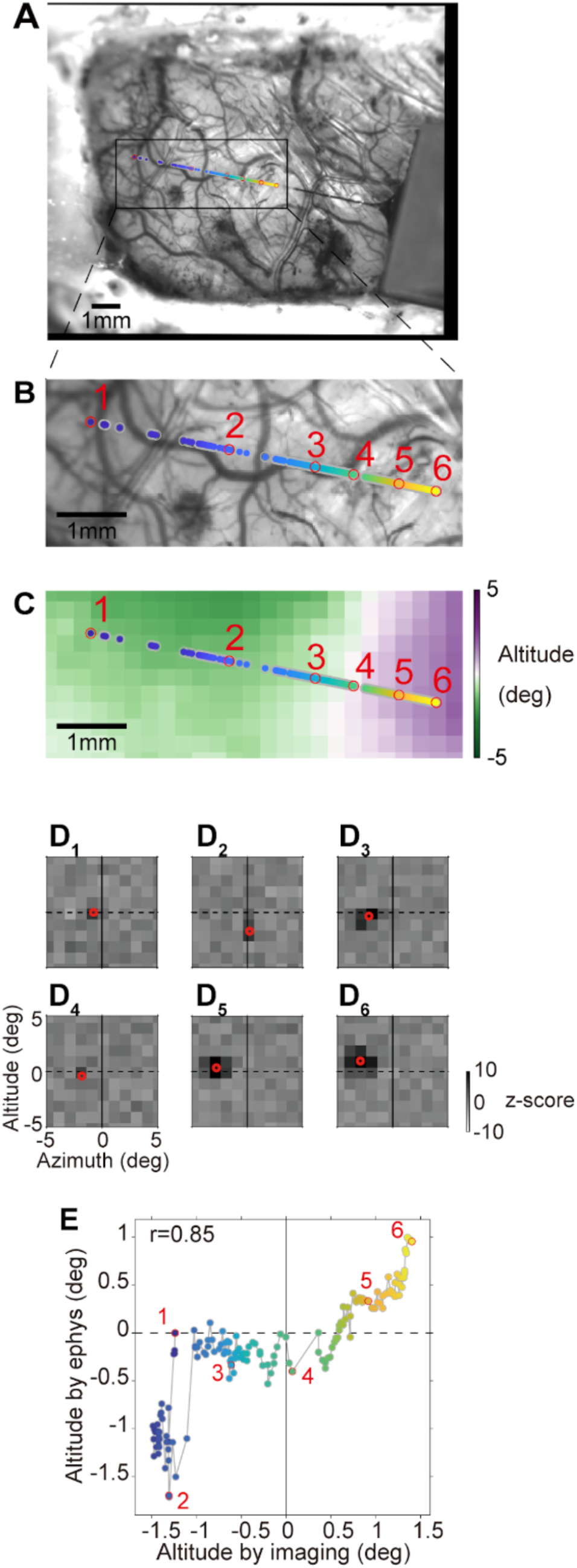
Relationship between Imaging and Electrophysiology Results for Preferred Visual Field Positions. **A.** Colored dots indicate locations of the electrophysiological recording channels with a significant number of detected spikes. The red circles indicate locations of the recording channels, used to compute receptive fields in panel **D**. **B.** Same image, enlarged around the locations of the recording channels. **C.** The locations of the recording channels are overlaid on the preferred visual field positions (altitude), obtained through imaging. **D.** Six receptive fields from different cortical locations are displayed. These receptive fields were computed based on the response to sparse noise stimuli. The center of the receptive fields (red circles) was obtained by fitting two dimensional gaussian curves. **E.** The preferred altitude in the visual field (abscissa), and electrophysiology (ordinate) from the same cortical locations, are compared. Notably, the preferred altitude estimated from both methods exhibit a high correlation (r=0.85), with both demonstrating a region that encodes the upper visual field (upper right quadrant).

### Visual response characterization using encoding model

Finally, we examined functional specialization in the form of speed tuning preferences in the area identified as DM. Speed is one of the fundamental visual properties of the motion processing, specialized in the dorsal stream in which DM and V3 are situated ^21^, and thus a candidate specialized property of these areas. From the encoding models fitted in each pixel, we obtained eccentricity of pRF (**Figure 5A**), preferred spatial frequency (**Figure 5B**), temporal frequency (**Figure 5C**) and speed (**Figure 5D**), defined by preferred spatial frequency divided by temporal frequency. Notably, the map of eccentricity exhibited a general correlation with the maps of spatial and temporal frequencies and speed exhibited, implying that as the receptive field position deviates from the visual field center or eccentricity increases, the preferred spatial frequency tends to decrease whereas the preferred temporal frequency and speed tend to elevate.

**Figure 5.**
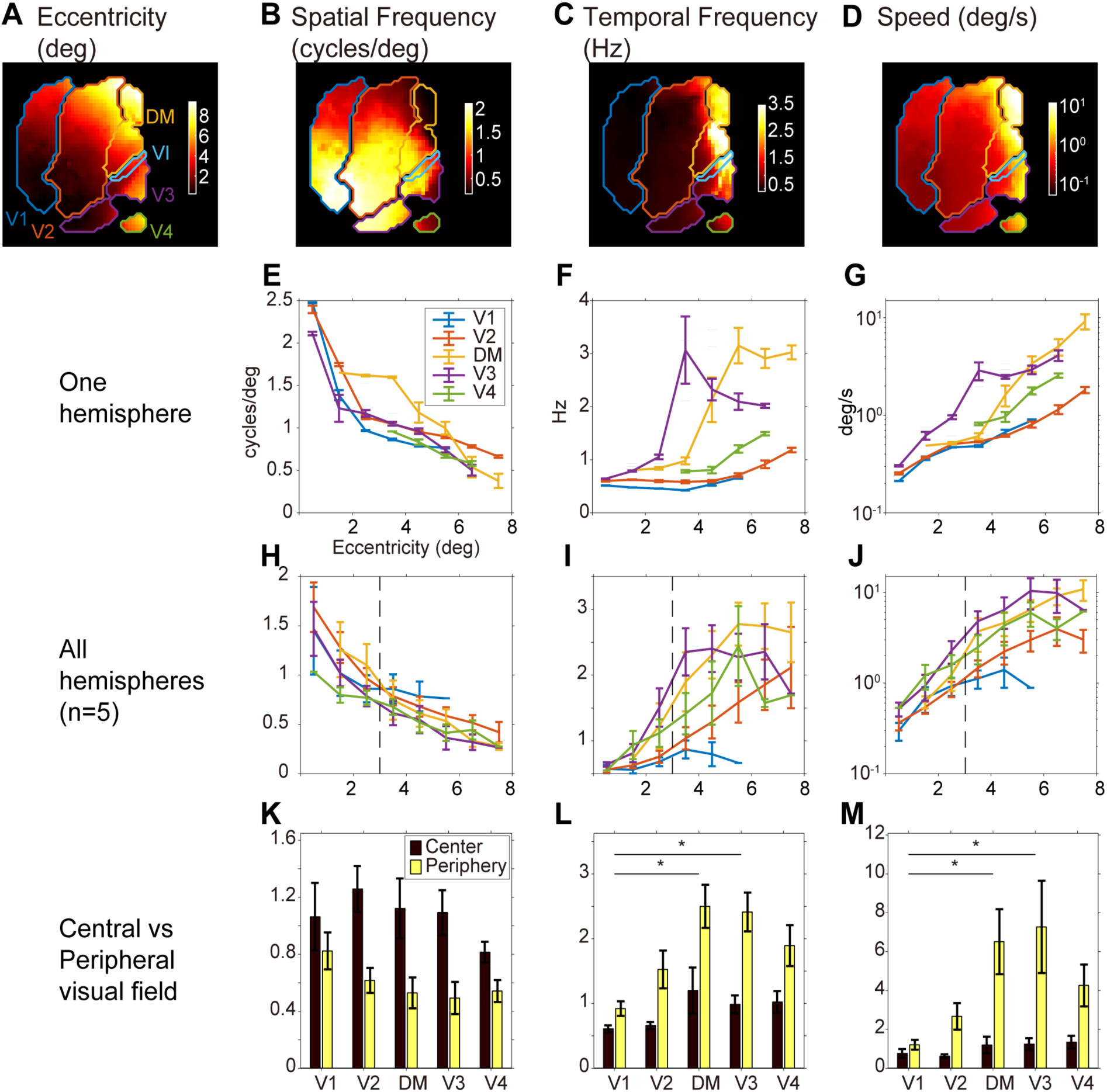
Spatio-temporal frequency and speed tuning characteristics across visual areas. **A**. Distance of receptive field position from the fovea in the visual field (eccentricity), in an example hemisphere. Colored contour lines represent borders of visual areas, determined from the visual field sign and the horizontal meridian (Fig. 3). **B-D**. Preferred spatial frequency, temporal frequency, and speed computed for each pixel in the same hemisphere, revealing a tendency for higher speeds in pixels with larger eccentricity. **E-G**. Preferred spatio-temporal frequency and speed, as a function of the eccentricity within the same hemisphere. Median and standard error of the median across pixels within each bin are presented. **H-J**. Preferred spatio-temporal frequency speed as a function of the eccentricity, averaged across the five hemispheres. **K-M**. Preferred spatio-temporal frequency and speed in the central (eccentricity < 3) and the peripheral (eccentricity >=3) visual field. In each area, the preferred speed is higher in the periphery. Mean and standard error of the mean across five hemispheres are presented. Notably, the preferred temporal frequency and speed in peripheral DM and V3 surpass those in peripheral V1.

To elucidate the eccentricity dependence of these response properties, we categorized pixels into bins based on eccentricity and their respective areal identities (**Figure 5E-G**). The preferred spatial frequency decreased at larger eccentricity in all five visual areas investigated. Likewise, the preferred temporal frequency and speed are clearly larger at higher eccentricities. Moreover, both DM and V3 displayed a marked inclination towards higher temporal frequency and speeds compared to the other areas investigated. These observations made in one hemisphere persisted across all the five hemispheres we investigated (**Figure 5H-J**).

To clarify the areal dependence of the preferred speed, we segregated pixels into central (less than 3 degrees eccentricity) and peripheral (3 degrees or more) visual fields, and evaluated the preferred speed across five hemispheres (**Figure 5K-M**). Intriguingly, the preferred temporal frequency and speed at the peripheral visual field of DM and V3 significantly surpassed that of V1. The difference could be due to the lack of observed pixels above 6 degrees eccentricity. However, we confirmed statistical difference when using pixels between 3 to 6 degrees. Together, these findings suggest a specialization of DM and V3 in encoding higher speeds.

## Discussion

### Summary of findings

In this study, we present a new methodology to delineate areal boundaries by integrating widefield imaging and encoding model analysis. Our approach resolves receptive field organization and stimulus tuning at a population level across a substantial portion of the cortex simultaneously. This results in an objective approach to defining areal borders, surpassing the capabilities of existing methods.

We applied this approach to investigate areal organization in and around the third-tier visual areas - whose precise organization remains elusive, particularly regarding how many areas form the dorsal visual cortex immediately rostral to V2, and where their boundaries lie ^14,15^. Across five hemispheres, our investigation consistently revealed: 1) a belt-like region at the anterior border of V2 representing the horizontal meridian; 2) a belt-like region representing the vertical meridian (at the border between V2 and V3); and 3) a region representing the lower visual field just anterior to a region of V2 representing the parafovea. Additionally, we identified a bulge region responsible for encoding the upper visual field, positioned immediately anterior to V2 and delineated by the horizontal meridian. This region was found to be medially and laterally adjacent to regions encoding the lower visual field. Finally, our approach further revealed the selective responsiveness of the third-tier areas in high temporal frequency and speed, in the peripheral visual field.

Our results represent a substantial advancement in understanding third-tier visual areas, particularly in contrast to previous imaging studies that yielded mixed outcomes. Although studies in galagos^22^ and owl monkeys^23–25^ successfully identified the rostral border of V2, regions rostral to V2 did not consistently exhibit reliable visual responses in previous functional imaging studies. Notably, compared to these studies employing predefined parameters of synthetic gratings, we adopted natural movies as visual stimuli. The natural movies contributed to enhancing the responsiveness of the third-tier visual areas to our stimuli, enabling acquisition of high resolution retinotopic maps and a comprehensive understanding of visual features encoded in each visual area.

### Case for multiple V3s

Our observations provide strong empirical evidence for the multiple third-tier areas hypothesis (**Figure 2B**). First, as we observed, the multiple V3’s hypothesis predicts the presence of a region encoding the upper visual field, located anterior and separated by V2 (**Figure 2C-E**). Second, we observed that the upper field-encoding region (**Figure 3**).

The existence of a narrow strip-like region rostral to V2, as suggested by previous studies, may provide evidence for the single third-tier area hypothesis^26–28^. This putative region is thought to be as slender as 2.8 mm in its width, based on observations in macaques. If a similar narrow strip were to exist in marmoset, its estimated width would be a mere 270 micrometers, inferred by the relative cortical surface sizes of macaque and marmoset^29,30^. It is possible that our imaging overlooked such a small structure. However, our electrophysiological recordings with a nominal resolution of 20 micrometers, did not reveal any region encoding the lower visual field between V2 and the region responsible for encoding the upper visual field. Consequently, our findings contradict the existence of this “narrow strip.”

A second argument supporting the single third-tier area pertains to the representation of the horizontal meridian within DM. It is suggested that DM displays a discontinuous representation of the horizontal meridian, featuring representations of the upper and lower visual fields with opposing field signs^31^. While such an organization might seem unconventional for a visual area, it has been theorized to jointly minimize wiring costs within DM and between DM and earlier visual areas, a proposition supported by electrophysiological evidence^11^. Our imaging results add further support to this notion, as they consistently reveal the upper and lower visual fields in the DM displaying opposing field signs across five hemispheres. Additionally, the observation of an S-shaped transition in the visual field (**Figure 2F**) is predicted by the elastic-net model explaining the formation of the opposing field signs.

Yet another possibility is that actual organization of the third-tier areas doesn’t align with either of these hypotheses. In line with this argument, Kaas^2,4^ has proposed an intriguing alternative possibility, suggesting that “dorsal V3 may have one or two gaps representing the upper visual field of smaller areas.” These gap regions were hypothesized to be spatially disconnected from the DM^2^. This idea gained some support from intrinsic imaging studies, reporting a representation of the lower visual field anterior to dorsal V2^22,32^. However, it is important to note that none of these studies reported any upper visual field representation adjacent to V2 or within the region of interest encompassing V2 and its surrounding areas, raising concerns about their validity. One plausible explanation for the lack of visual response from the third-tier areas in this imaging study could be the use of moving gratings with fixed spatio-temporal frequencies, which may not have been optimal for activating all regions of the third-tier visual areas, presumed to cover a wide range of eccentricities and spatial frequencies. Furthermore, stimulus-evoked responses are inherently susceptible to artifacts caused by blood vessels, complicating the interpretation of whether the apparent “patchiness” of visually-responsive regions^16^ is due to this artifact or the columnar organization of the visual area.

In our approach, we mitigated the potential for arbitrary stimulus parameter selection by combining natural movies with encoding modeling. Moreover, our method is less vulnerable to the artifact caused by blood vessels, partly due to the enhanced depth of field achieved with light-polarization techniques. Using this approach, we discovered a region encoding the upper visual field. Unlike the “gap” hypothesis, this region was found to be connected to the DM, which encodes the lower visual field. Consequently, in terms of functional retinotopy, our findings align more with the multiple-V3 hypothesis, rather than the “gap” proposal.

Together, our observations provide strong empirical evidence for the multiple third-tier areas hypothesis.

### Utility of encoding model approach

Encoding modeling has been recently used to explore encoding of abstract modalities such as emotion (e.g. ^33^) and cognition (e.g. ^34^) in fMRI signals. Our approach, which combines encoding models and widefield optical imaging, holds the potential to contribute to the resolution of other contentious cortical areas, such as the Posterior Parietal Cortex, by incorporating modalities other than vision such as bodily motion into the encoding model^35–38^. This methodology not only aids in the delineation of areal boundaries but also provides the means to characterize aspects of visual signals each area processes. As an example, this study demonstrated distinct processing of higher temporal frequency and speed in the third-tier visual areas. The tuning in higher speed in third-tier areas is also reported in previous studies (Marmoset: ^21^; Human: ^13^). In future studies for exploration of even higher visual areas, it is essential to employ quantitative models of neuronal responses specifically tailored for higher visual areas. Through these processes, future research endeavors will have the opportunity to uncover the distinctive neural response characteristics inherent to each area, thus furthering our understanding of their roles as modules in visual information processing.

## Methods

All procedures were approved by the Monash University Animal Ethics Committee and were conducted in compliance with the Australian Code for the Care and Use of Animals for Scientific Purposes.

### Surgical procedure

Two male and two female marmoset monkeys (C. jacchus) were used. Animals were first given intramuscular atropine (Atrosite, 0.33ml/kg) and diazepam (Pamlin, 0.4ml/kg), then the anesthetic alfaxalone (Alfaxan, 8 mg/kg). The animals received an antibiotic (Duplocillin, 25.5 mg/kg) and dexamethasone (Dexason, 0.3 mg/kg). A tracheotomy was performed, and the femoral vein cannulated. The monkeys were positioned in a stereotaxic frame with a head-fixation bar, and a craniotomy was performed over the dorsomedial extrastriate portion of the cerebral cortex (from 11 mm posterior to 2 mm anterior to the external auditory meatus) with a dental drill (Volvere i7, NSK, Japan). The exposed cranial window (**Figure 1C**), covered an area of approximately 10×13 mm, designed to encompass multiple visual areas including V1, V2, third-tier-visual areas and posterior-parietal cortex, in accordance with a published atlas (Liu et al., 2021).

Five hemispheres (4 from right, 1 from left hemispheres) from four animals (Hemispheres B and C from the same animal) were subsequently recorded. For visualization purpose, the result from the left hemisphere (Hemisphere C), was mirror transformed. During the recording session, the monkeys were maintained on an infusion of Sufentanil citrate (Sufenta Forte, 250 ug/5 ml), Vecuronium (4 mg/2 ml), dexamethasone (Dexapent, 5 mg/ml), xylazine (Xylazil-20, 20 mg/ml), and a solution containing salts and nutrients (0.18% NaCl/4% glucose solution, Synthamine-13, and Hartmann’s solution). They were ventilated using a mixture of nitrous oxide and oxygen in a 7:3 ratio. The monkeys’ body temperature was maintained at a constant 38°C, monitored with a rectal thermometer.

The contralateral eye to the craniotomy was held open, and before inserting a contact lens to focus the eye at a viewing distance of 50-80 cm, atropine (Atropt, 1%), phenylephrine, and carmellose sodium (Celluvisc) eye drops were applied. The ipsilateral eye was protected with carmellose sodium, closed, and occluded.

### Visual stimulation

Visual stimuli were presented on a gray background using a liquid crystal display (Display++, Cambridge Research Systems, UK) positioned at a viewing distance of 50-80 cm. The Psychophysics Toolbox in MATLAB, in conjunction with its wrapper neurostim (http://klabhub.github.io/neurostim/), was used to design and present the visual stimuli at a refresh rate of 120 Hz.

The pseudo-white noise stimulus consisted of squares measuring 0.5°-1°, alternating between “on” (white) and “off” (black), and were presented for 50 ms on a gray background. Each square appeared for 50-100 ms (varying depending on the specific case).

The natural movie stimuli comprised short clips lasting approximately 10 seconds each. These clips depicted various natural scenes, including moving animals, humans, letters, and sentences ^13^. The movie stimuli were obtained from the publicly available dataset VIM-2 (https://crcns.org/data-sets/vc/vim-2/about-vim-2) and were presented at a frame rate of 15 Hz. In total, twenty-four 5-minute movies were shown, resulting in a cumulative duration of 120 minutes, with 10 seconds between individual short clips.

### Widefield intrinsic optical imaging

The cranial window was contained mechanically in the following procedure. Around the cranial window was secured by a 3D-printed cone with superglue (zap-a-gap, Pacer Technology, USA). It was then covered with transparent agar (cat no 800257, MP biomedicals, USA) and fused by a 3D-printed lid housing a glass coverslip, attached to the wall of the cone. This way of containment enabled stable imaging by suppressing the pulsation of the brain and preventing the ambient light entering the optical path for the imaging.

Widefield imaging was conducted with an optical system based on tandem-lens design (Imager 3001, Optical Imaging, Israel), consisting of two macro lenses. One lens (SDFPLAPO 0.5XPF, Olympus, Japan) faced the contained cranial window whereas the other lens (50 mm F/1.2, Nikon, Japan) faced a CCD camera (1000m, Adimec, Netherland). We recorded the “intrinsic” signal of the cortex, originating from the blood flow change incurred by the neuronal spiking activity as follows: The contained cranial window was exposed to green (530 nm) light (M530L4, Thorlabs, USA), and reflected light was collected with the camera recorded at 10 Hz. A linear polarization filter was placed in front of the light source (WP25L-VIS, Thorlabs, USA), and a second polarization filter, orthogonal to the first, was placed within the collection light path. As a result, the camera recorded “cross-linear” photons that have lost their initial polarization and tend to reach deeper brain structures ^39^.

### Electrophysiological recording

In a subset of experiments, widefield imaging was followed by electrophysiological recording from the same cranial window. A small cut was made in the dura and a neuropixels 1.0 probe was inserted, as close to tangential to the cortical surface as possible. The cranial window was then covered with agar. The probe was advanced at 4 um/s to a depth of 7.7 mm, and then left still for at least 30 minutes before the start of data acquisition. Electrophysiological signals were acquired and monitored using openEphys GUI 0.5 (https://open-ephys.github.io/). Signals from 384 channels, separated by 20 um, were recorded with a custom channel mapping file.

### Signal processing

#### Encoding-model analysis

Imaging signal during exposure to natural movies was first down-sampled spatially, resulting in a nominal pixel size of 400×400 um. In the temporal domain, the imaging signal underwent high pass filtering above 0.02 Hz, followed by down sampling at 1 Hz.

The natural movie was transformed into monochrome (CIE 1976 L*a*b*) then convolved with a bank of Gabor-Wavelet filters of the motion-energy model^13^. The model operates through a series of steps to process visual stimuli. Initially, the monochrome signals are passed through a bank of 6,555 spatiotemporal Gabor filters, with defined positions, orientations, directions, spatial frequencies, and temporal frequencies. The filters are constructed by multiplication of a three-dimensional spatiotemporal sinusoid by a three-dimensional spatiotemporal Gaussian envelope. These filters are distributed across the visual field, with varying spatial frequencies (ranging from 0 to 32 cycles per image), temporal frequencies (0, 2, and 4 Hz), and orientations (0, 45, 90, and 135 degrees) to comprehensively capture motion characteristics. Notably, filters associated with zero temporal frequency are limited to only four orientations, while those with zero spatial frequency occur just once without any specific orientation.

To ensure an effective coverage of the visual stimuli, these filters are arranged in a grid pattern that spans the entire screen. The spacing between adjacent Gabor wavelets is determined individually for each spatial frequency, guaranteeing that they are separated by 3.5 standard deviations of the spatial Gaussian envelope.

Following processing through these Gabor filters, the signals are log transformed then temporally downsampled to a rate of 1 Hz and converted to z-scores.

The z-scored output of the motion-energy model was used to fit the down-sampled imaging signal for each camera pixel with L2-regularized linear regression. The linear regression coefficients correspond to the hemodynamic coupling term in the motion-energy model. To minimize computational time, we restricted the temporal window of the hemodynamic coupling terms to a period of 2-4 seconds (3 time points) before imaging signals, without using the period of 0-1 seconds. After successfully fitting the motion-energy model, our next step was to estimate its receptive field. Visualizing this receptive field was challenging due to the characteristic of the motion-energy model, which comprises numerous Gabor wavelets positioned at various locations and scales. Additionally, the model incorporates hemodynamic delays that are distinct for each motion-energy filter and pixel, further complicating the visualization process. We estimated spatial receptive fields via *in silico* simulation, where we simulated the model response to a set of sparse noise stimuli. The simulated response after 2-4 s of each noise patch was averaged, and its response peak was obtained by fitting a two-dimensional Gaussian in the visual field domain. The resulting peak location was registered as the preferred position of a population receptive field (red circle in **Figure 1F**) for each camera pixel. By performing this procedure across all camera pixels, the preferred positions were identified in relation to the foveal representation. The latter was obtained from the most lateral-posterior pixel within the cranial window, and defined as (azimuth, altitude) = (0, 0), corresponding to the foveal representation of the primary visual cortex. The resulting preferred positions were visualized as maps of preferred azimuth and altitude on the cortical surface, or maps of distance from the foveal representation (eccentricity). Notably, these maps were shown as they were, without any spatial smoothing or filtering across pixels. We determined if the motion-energy model sufficiently explained the observed signal by measuring correlation between the observed and modeled signals. Only the regions with high correlation values (0.24-0.29 depending on hemispheres) were represented.

#### Detection of areal boundaries

After establishing the preferred positions within the visual field for all the camera pixels, the visual field sign was determined by computing the chirality of these preferred positions across the cortical surface^40^. More specifically, the chirality was computed by determining the gradient within the preferred altitude and azimuth maps and taking the sine of the difference between the azimuth and elevation gradients. The visual field sign was then leveraged to derive estimates of boundaries between distinct visual areas. The field sign map from individual hemispheres were averaged through an image translation (**Figure 3B**). This transformation allowed for rotation and translation adjustments (rigid registration) while minimizing the mean squared error, achieved through a gradient descent algorithm (Mean Squared Error across 3 pyramid levels: 0.13, 0.20, 0.33 (Hemisphere B); 0.14, 0.23, 0.37 (Hemisphere C), 0.09, 0.14, 0.26 (Hemisphere D), 0.09 0.16, 0.26 (Hemisphere E). Additionally, building upon the finding that horizontal meridian can serve as a delineating factor for areal borders ^11^, a line representing horizontal meridian on the cortex was estimated from the cortical map of the preferred position.

#### Electrophysiological data analysis

During the acquisition of electrophysiological data, a series of sparse noise stimuli were presented over a duration of approximately 30 minutes. The spike times of each recording channel were detected based on adaptively set thresholds^41^. Units with more than 1000 detected spikes during the stimulation period were processed for further analysis. To derive the receptive field associated with each recording channel, stimulus-triggered average response was calculated following a time interval of 40-130 ms. The center of the receptive field was obtained by least-square fitting a two-dimensional Gaussian curve to the stimulus-triggered average response in the visual field domain. Units with poor fitting were excluded from further analysis (detected as the residual of fitting, normalized by the number of the stimulation squares to be larger than 0.013). Units with a small response amplitude, detected as peak amplitude / baseline to be less than 0.7, were also excluded. The receptive field positions in the visual field were translated according to the preferred visual field position at the posterior-lateral region with widefield imaging.

#### Alignment of electrophysiological recording into widefield imaging

We took a picture of the brain during the electrophysiological recording, with the same camera at the same angle as those during widefield imaging. The picture was later aligned to the image during widefield imaging using non-reflective similarity transformation. Using this aligned image, we estimated the location of the recording channels projected onto the cortical surface.

#### Speed tuning across different areas

Utilizing the encoding models obtained through fitting, we examined their preferred speed via *in silico* simulation. In this simulation, we replicated the model’s response to a range of drifting-grating stimuli characterized by different directions and spatio-temporal frequencies. The accumulated response during the 2-4 seconds following the onset of each stimulus was averaged and depicted as two-dimensional representation as a function of spatial and temporal frequencies. The preferred temporal and spatial frequencies were obtained by fitting a two-dimensional Gaussian. The preferred speed was subsequently derived by dividing the preferred temporal frequency by the preferred spatial frequency.

To assess the areal dependence of the preferred spatio-temporal frequencies and speed, we aggregated pixels into distinct visual areas based on the visual field sign and the horizontal meridian. In each area, pixels with a metric value exceeding 6 times the median absolute deviation were excluded. This removed 0.4% of data for each area across hemispheres. To analyze eccentricity dependence of the preferred speed, pixels were grouped into equally spaced bins ranging from 0 to 8 deg eccentricity. The eccentricity dependence of the preferred speed was computed through a robust regression in each hemisphere, followed by a statistically testing of their areal difference using the Kruskal-Wallis one-way analysis of variance. Additionally, bins were combined into central (<3 deg eccentricity) and peripheral (>= 3deg) visual fields. With each category, the average preferred speed in each subject was obtained, and their areal difference were statistically tested using the Kruskal-Wallis test. Further testing of areal difference of pixels between 3-6degrees, Wilcoxon-sign rank test was performed between each pair of areas.

## Acknowledgements

Authors would like to thank David Reser for providing the materials necessary for conducting surgical procedures, Elizabeth Zavitz for feedback on earlier version of the manuscript, and all members of the lab for supporting the experiments conducted.

